# Crocin inhibited amyloid-beta (Aβ) generation via promoting non-amyloidogenic APP processing and suppressed ER stress UPR signaling in N2a/APP cells

**DOI:** 10.1101/2021.09.14.460251

**Authors:** Crystal Cuijun Lin, Erin Qian Yue, Zirong Liang, Simon Ming Yuen Lee, Zaijun Zhang, Maggie Pui Man Hoi

**Author notes:** **Corresponding Author:** Dr Maggie Pui Man Hoi, Institute of Chinese Medical Sciences, University of Macau, N22-7012, Avenida da Universidade, Taipa, Macau, China.

## Abstract

**Background:** Crocin is a major active component of saffron (*Crocus sativus*) with many beneficial effects. More recently, crocin has been proposed for management of neurodegenerative diseases such as Alzheimer’s disease (AD). Here, we demonstrated for the first time that crocin reduced amyloid-beta (Aβ) generation through promoting α-cleavage of APP processing and inhibited ER stress by attenuating UPR signaling in N2a/APP cells.

**Methodology:** Mouse neuroblastoma N2a cells stably transfected with the Swedish mutant APP (N2a/APP) was used as a cellular model for AD pathogenesis. Vector transfected cells (N2a/vector) were employed to serve as control. The toxicity of crocin was first evaluated and non-toxic treatment of crocin (>30 µM for 24 h) was used for further investigations. Aβ levels were determined by ELISA. Expression levels of UPR signaling proteins were determined by using Western blot.

**Results:** Crocin significantly inhibited the protein expression of total APP in N2a/APP cells and promoted α-cleavage of APP processing to increase sAPPα generation, but only modestly reduced BACE-1 and PS1, suggesting Aβ reduction by crocin was mainly associated with the non-amyloidogenic APP processing. Further investigation on ER stress related protein expressions showed that GRP78, CHOP, p-PERK, p-eIF2α, p-IRE1α, XBP1, ATF6α and PDI were all significantly elevated in N2a/APP cells compared to N2a/vector. Crocin effectively reduced the levels of GRP78 and CHOP, and significantly inhibited p-PERK/p-eIF2α amd AT6α while slightly reduced p-IRE1α.

**Conclusion:** The present study showed that crocin was effective at blocking Aβ generation and inhibiting ER stress associated overactivation of UPR signaling in AD cell model N2a/APP. The results provided evidence for crocin as useful natural product for the treatment of AD.

## Introduction

Crocin (trans-crocetin di-β-D-gentiobiosyl ester, C_44_ H_64_ O_24_, Fig. 1A) (also known as crocin-I or α-crocin) is a water-soluble carotenoid and a major active component of saffron (*Crocus sativus*) [1]. Crocin possesses many therapeutic effects due to its strong antioxidant, anti-apoptotic and anti-inflammatory properties. More recently, crocin has been proposed for management of neurodegenerative diseases such as Alzheimer’s disease (AD). Studies of crocin in AD-related mouse and rat models have provided strong evidence for their beneficial effects against cognitive decline and neuronal cell death. In a study using transgenic mouse model 5xFAD [2], crocin was demonstrated to increased BBB functions and reduced brain amyloid bet (Aβ) load by upregulating Aβ clearance across the BBB, increasing expression of Aβ degrading enzyme NEP, increasing ApoE clearance and reducing neuroinflammation associated with Aβ pathology. In a AD-like mouse model induced by D-galactose and aluminum trichloride [3], crocin significantly improved memory performance and reduced Aβ levels in the brain homogenates, increased defense antioxidants such as glutathione peroxidase (GPx) and superoxide dismutase (SOD) and alleviated cholinergic dysfunction. In various AD-related rat models, crocin (intra-peritoneal, intra-hippocampal or oral administration) improved cognitive functions and memory performance, prevented the elevation of oxidative stress and the accumulation Aβ and hyperphosphorylated tau (pTau), and attenuated Aβ-induced cellular apoptosis in the brain [4-7].

**Fig. 1.**
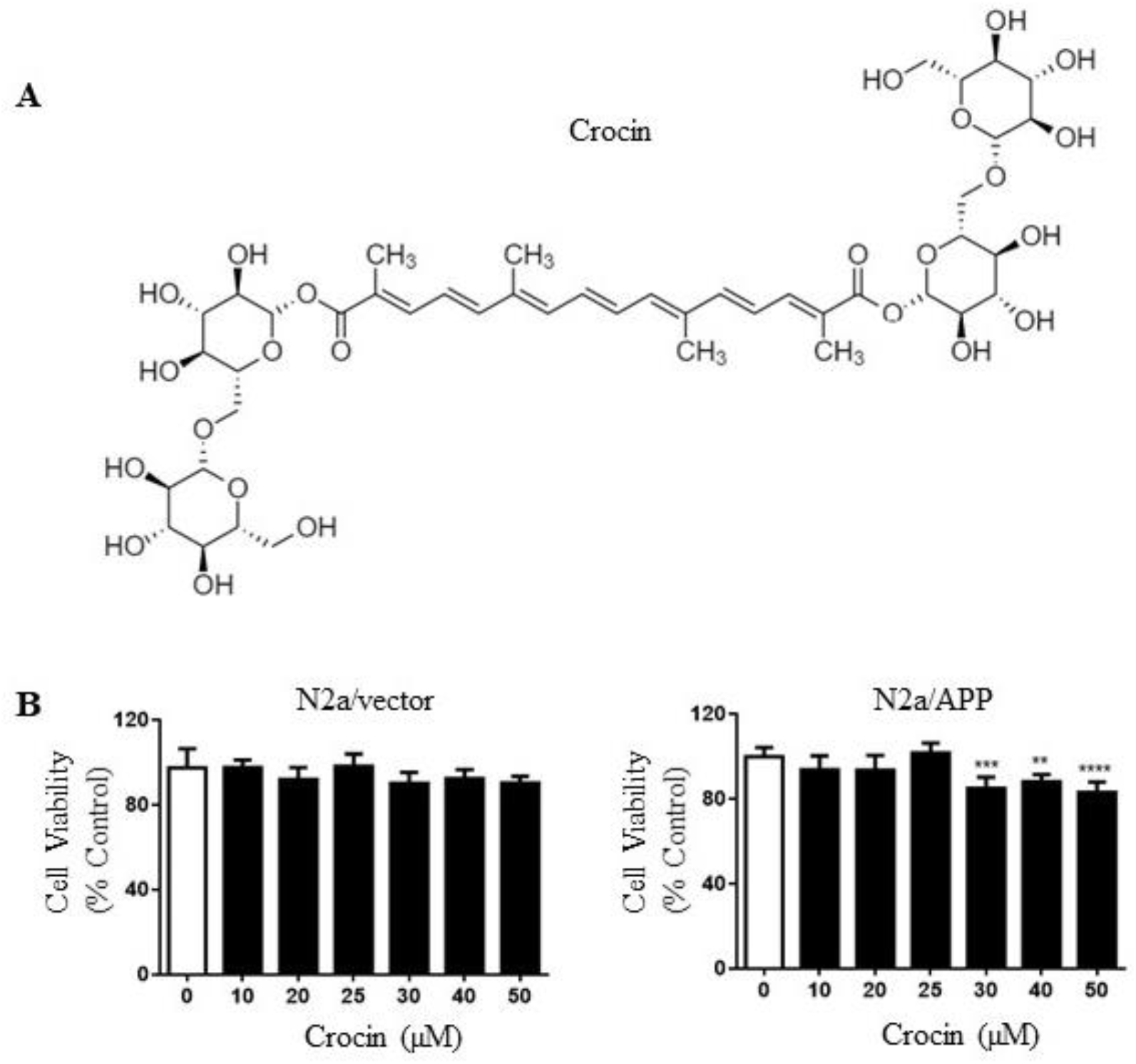
(A) Chemical structure of crocin. (B) Cell viability of N2a/Vector and N2a/APP cells after crocin treatment (24 h). n = 5. \*\**p* < 0.01, \*\*\**p* < 0.001,\*\*\*\**p* < 0.0001 compared with N2a/APP cells treated with vehicle.

The aggregation of misfolded proteins occurs in many neurodegenerative diseases, such as in AD the accumulation of abnormally folded Aβ and pTau in amyloid plaques and neurofibrillary tangles are the clinical hallmarks of the disease [8]. Both familial and sporadic forms of AD have similar Aβ and pTau pathologies but with different time of onset, suggesting common pathogenic mechanisms. Familial AD is caused by mutations in *APP, PSEN1* and *PSEN2*, which are the genes associated with Aβ generation [9]. The amyloid precursor protein (APP) can undergo sequential proteolytic processing via the non-amyloidogenic (α-secretase pathway) and the amyloidogenic (β-secretase pathway). In the non-amyloidogenic pathway, α-secretase cleaves within the Aβ peptide sequence region of APP to produce sAPPα and the membrane-tethered C83, which is subsequently processed by γ-secretase to produce p3 and AICD. In the amyloidogenic pathway, alternative cleavage of APP by β-secretase results in the generation and secretion of sAPPβ and the membrane-tethered C99, which is subsequently processed by γ-secretase to produce Aβ And AICD [10, 11]. Therefore, the upregulation of APP and dysregulation of APP processing play pivotal roles in AD pathogenesis.

The endoplasmic reticulum (ER) is a major protein folding compartment for newly-synthesized proteins and is one of the primary intracellular sites in which misfolded proteins is accumulated. Improperly folded proteins are retained in the ER by resident chaperone as quality control mechanisms. When the rate of protein synthesis exceeds the capacity of the ER to direct proper protein folding, elevated levels of unfolded proteins results in ER stress that initiates response mechanisms known as the unfolded protein response (UPR). The primary aim of UPR signaling is to restore ER homeostasis by halting protein translation, degrading misfolded proteins and increasing production of chaperones involved in protein folding. However, sustained UPR overactivation triggers apoptotic signaling during prolonged stress conditions. In AD patients, especially within the pathologically affected brain regions, there is overactivation of UPR signaling and suggests ER stress and UPR contribute to the pathological progression in AD [12].

The present study aims to evaluate the beneficial effects of crocin for treating AD. We investigated the effects of crocin on Aβ production, APP processing and UPR signaling by using mouse neuroblastoma N2a cells stably transfected with the Swedish mutant APP (N2a/APP). Vector transfected cells (N2a/vector) were used to serve as control.

## Methods

### Chemicals and Reagents

Crocin (purity > 98%) was purchased from Sichuan Victory Biological Technology Co., Ltd. (Sichuan, China). Dulbecco’s Modified Eagle Medium (DMEM), fetal bovine serum (FBS), penicillin-streptomycin (P/S), phosphate-buffered saline (PBS), 0.25% trypsin-EDTA were purchased from Gibco/Invitrogen (USA). Dimethyl sulfoxide(DMSO), MTT[3-(4,5-dimethylthiazol-2-yl)-2,5-diphenyltetrazoliumbromide] were acquired from Sigma (USA). Purified water was prepared with a Milli-Q purification system from Millipore (Milford, USA). Human Aβ (aa1-40)/(aa1-42) quantikine ELISA Kits were purchased from R & D systems (Minneapolis, MN, USA). CM-H2DCFDA (General Oxidative Stress Indicator), MitoTracker Red CM-H2XRos were purchased from Thermo Fisher Scientific (NJ, USA). β-actin antibody (Cat. No: 4970S), GAPDH antibody (Cat. No: 2118), Anti-rabbit IgG, HRP-linked antibody (Cat. No: 7074S), Anti-mouse IgG, HRP-linked antibody (Cat. No: 7076S), Bip/Grp78 antibody (Cat. No: 3183S), p-elf2α antibody (Cat. No: 9721S), elf2α antibody (Cat. No: 9722S), presenilinl (D39D1) antibody (Cat. No: 5643S) were all purchased from Cell Signaling Technology (Berverly, MA). Chop antibody (Cat. No: 15204-1-AP) were all purchased from Proteintech (USA). EIF2AK3 (LS-C118320) antibody (Cat. No: 38117), EIF2AK3 (LS-C199431) antibody (Cat. No: 49990) were all purchased from LSBio (USA). ATF6 antibody (Cat. No: SC22799) were purchased from Santa Cruz Biotechnology (Dallas, TX, USA). IRE1 (phospho S724) antibody (Cat. No: ab48187), IRE1 antibody (Cat. No: ab37073), XBP1 antibody (Cat. No: ab37152), Amyloid Precursor Protein antibody antibody (Cat. No: ab15272), BACE1 antibody (Cat. No: ab2077) were purchased from Abcam (Cambridge, MA, USA). Antibody sAPPβ were purchased from IBL (Minneapolis, MN, USA).

### Cell Culture

Mouse neuroblastoma Neuron2a cells stably transfected with human APP Swedish mutant (N2a/APP) were a gift from Dr. Gopal Thinakaran (University of Chicago, Chicago, IL, USA) and obtained from Dr. Jiahong Lu (University of Macau, Macau, China S.A.R. [13]). Neuron2a cells transfected with empty vector (N2a/vector) were used as control. The cells were cultured in DMEM contain with 10% FBS (Gibco), 100 U/ml penicillin (Gibco), 100 μg/ml streptomycin and 200 μg/ml G418 (Solarbio). Cells were incubated at 37°C in 5% CO_2_ :95% air environment.

### Cell Viability MTT Assay

N2a/vector or N2a/APP cells were trypsinized and seeded at 1.5×104/cm^2^ in 96-well plates. Cells were treated with crocin for 24 h. Then the medium was discarded and cells were incubated for 4 h at 37°C in 0.5 mg/ml MTT solution. The solution was then replaced by 100 μl DMSO to dissolve the violet formazan crystals in intact cells. The absorbance was measured at a wavelength of 570 nm by a SpectraMaxR M5 Multi-Mode Microscope Readers. Cell viability was expressed as a percentage of the vehicle control.

### Determination of Aβ_1-40_ and Aβ_1-42_ contents by ELISA

The levels of secreted Aβ contents were determined by using a sandwich ELISA as previously described [13]. Briefly, cell culture medium was collected and cleared by centrifugation for use in detecting extracellular Aβ levels. The levels of Aβ_1-40_ and Aβ_1-42_ were quantified by Quantikine ELISA Human Amyloid β (aa1-40/aa1-42) Immunoassay kits following the manufacturer’s instructions.

### Western Blot

Protein samples were extracted from cells using ice-cold RIPA buffer with complete protease inhibitor mixture (Roche Applied Science, 04693124001). For immunoprecipitation, cells were lysed in cell lysis buffer (25 mM Tris, pH 7.6, 100 mM NaCl, 0.5% NP40 (Sigma), 1 mM EDTA, 10% glycerol with protease inhibitors. After immunoprecipitation with the indicated antibodies, proteins were resolved by gel electrophoresis in 10–15% SDS-polyacrylamide gels, and subsequently transferred onto PVDF membranes. Following blocking with TBS-T (Tris-buffered saline with 0.1% Tween-20 (Sigma) buffer containing 5% nonfat milk powder, the blots were probed with specific primary antibodies, and finally visualized using Pierce ECL kit (Thermo Scientific Pierce ECL, USA). Quantitative densitometry analyses were performed using Quantity One Analysis Software.

### Statistical Analysis

All data were expressed as mean ±standard error of the mean (S.E.M.) of at least three independent experiments. Differences between means were evaluated with one-way ANOVA for comparison of three or more groups. .*p* < 0.05 was considered to be statistically significant. \**p* < 0.05, \*\**p* < 0.01, \*\*\**p* < 0.001, \*\*\*\**p* < 0.0001.

## Results

### 1. Crocin has low cytotoxicity for N2a/vector and N2a/APP cells

We first studied the cytotoxicity of crocin on N2a/vector and N2a/APP cells. Crocin (1-50 µM for 24 h) did not induce any decrease in cell viability in in N2a/vector cells. For N2a/APP cells, crocin slightly reduced cell viability at concentrations > 30 µM (Fig. 1B). Accordingly, crocin treatment with a maximum concentration of 25 µM for 24 h was used in the following investigations.

### 2. Crocin reduced Aβ production by regulating APP processing pathway in N2a/APP cells

In order to study the effect of crocin on Aβ production, we determined the levels of Aβ_1-42_ and Aβ_1-40_ in the culture media of N2a/vector and N2a/APP cells with or without crocin treatment. As shown in Fig. 2, the level of “toxic” Aβ_1-42_ and “non-toxic” Aβ_1-40_ in the culture medium of N2a/APP were much higher than that in N2a/vector. The ratio of Aβ_1-42_ /Aβ_1-40_ was also significantly increased. The treatment of crocin (≤ 25 µM for 24 h) significantly reduced the level of Aβ_1-42_ released by N2a/APP (Fig. 2A) and reduced the level of Aβ_1-40_ more modestly (Fig. 2B). The ratio of Aβ_1-42_ /Aβ_1-40_ was also reduced by crocin (Fig. 2C). We further determined the expression levels of the proteins involved in the APP processing pathway, including t-APP, sAPPα, sAPPβ, BACE1 and PS1 (Fig. 3A). The level of t-APP protein was significantly reduced by crocin (Fig. 3B). Interestingly, the level of sAPPα was significantly increased by crocin (Fig. 3C). The levels of sAPPβ, BACE1 (β-secretase) and PS1 (γ-secretase) were moderately reduced by higher concentration (25 µM) of crocin (Fig. 3D-F). These results suggested that crocin reduced Aβ production by reducing total APP production as well as upregulating the cleavage of APP by α-secretase to generate sAPPα, therefore reducing sAPPβ and downstream Aβ production.

**Fig. 2.**
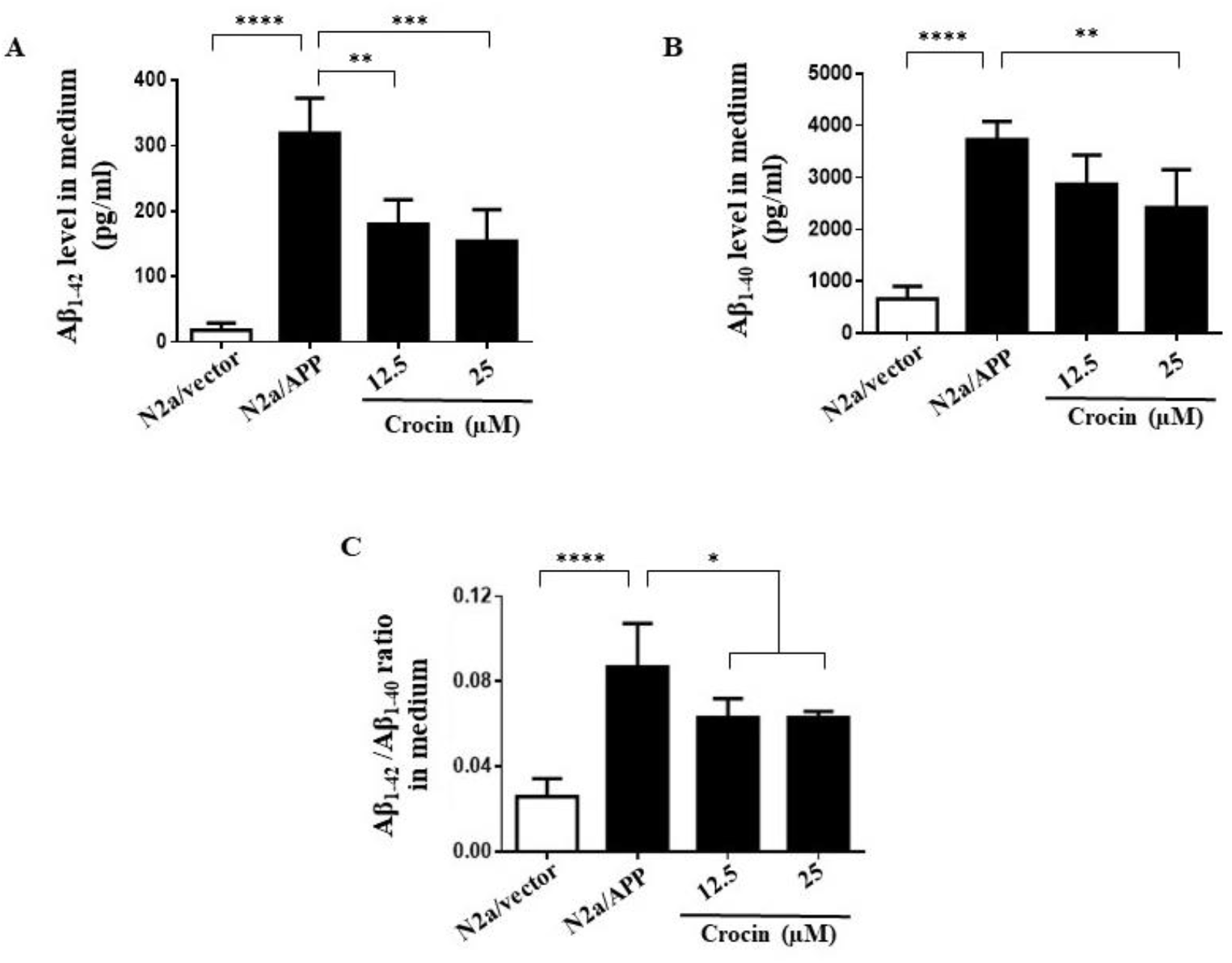
The production of Aβ_1-42_ and Aβ_1-40_ were significantly increased in N2a/APP cells and crocin treatment (≤ 25 µM for 24 could inhibit it. (A,B) Crocin reduced Aβ_1-42_ level potently and reduced Aβ_1-40_ level more modestly. (C) The ratio of Aβ_42_ /Aβ_40_ was also reduced by crocin treatment. n =3. \*\**p* < 0.01, \*\*\**p* < 0.001,\*\*\*\**p* < 0.0001 compared with N2a/APP cells treated with vehicle.

**Fig. 3.**
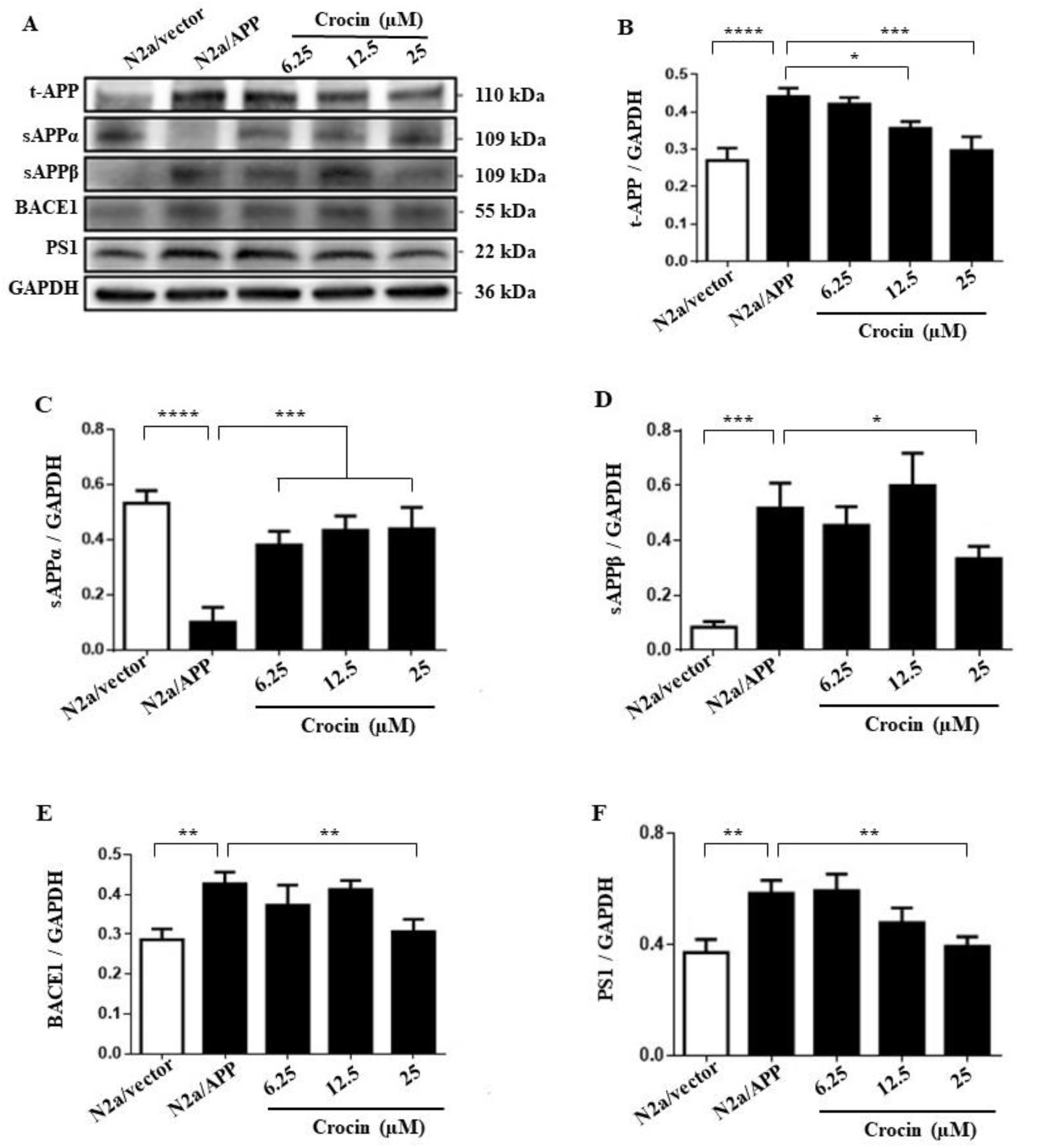
The treatment of crocin (24 reduced Aβ production in N2a/APP cells by regulating the APP processing pathways. (A) Western blots of proteins in APP processing pathways. (B-C) The level of t-APP was significantly reduced by crocin, while the level of sAPPα was significantly increased. (D-F) The levels of sAPPβ, BACE1 and PS1 were moderately reduced by crocin. n = 3. \*\**p* < 0.01, \*\*\**p* < 0.001,\*\*\*\**p* < 0.0001 compared with N2a/APP cells treated with vehicle.

### 3. Crocin attenuated the upregulation of ER stress and UPR signaling in N2a/APP cells

ER stress is implicated in AD pathogenesis. The accumulation of misfolded proteins and perturbation of intracellular Ca^2+^ homeostasis are thought to underlie the induction of ER stress resulting in neuronal dysfunction and cell death. ER stress activates the adaptive unfolded protein response (UPR) signaling pathways that consist of three key branches, IRE1α/XBP1s, PERK/eIF2α/ATF4, and ATF6α. GRP78 is a master senor to initiate UPR, and CHOP is a transcription factor that is activated at multiple levels during ER stress. Here we observed that ER stress was induced and UPR signaling was significantly upregulated in N2a/APP cells, as indicated by the upregulation of GRP78, CHOP, p-PERK, p-eIF2α, p-IRE1α, XBP1, and ATF6α (Fig. 4A-G). Under normal conditions, GRP78 is localized to the ER lumen, and PERK, IRE1α, and ATF6α remain in inactive state. Upon ER stress, misfolded proteins inhibit interaction between GRP78 and initiates UPR signaling. The treatment of crocin significantly attenuated GRP78 and CHOP expressions in N2a/APP cells (Fig. 4A-B). Crocin effectively inhibited the activation of p-PERK and p-eIF2α (Fig. 4C-D) and the expression of ATF6α (Fig. 4G). The effect of crocin on the activation of p-IRE1α was more modest (Fig. 4E) while crocin was able to significantly reduce the protein level of total XBP1 (Fig. 4F). The protein level of PDI, which is a key player in ER homeostasis with protein folding and chaperone properties, was increased in N2a/APP cells and crocin did not have any effect on PDI (Fig. 4H).

**Fig. 4.**
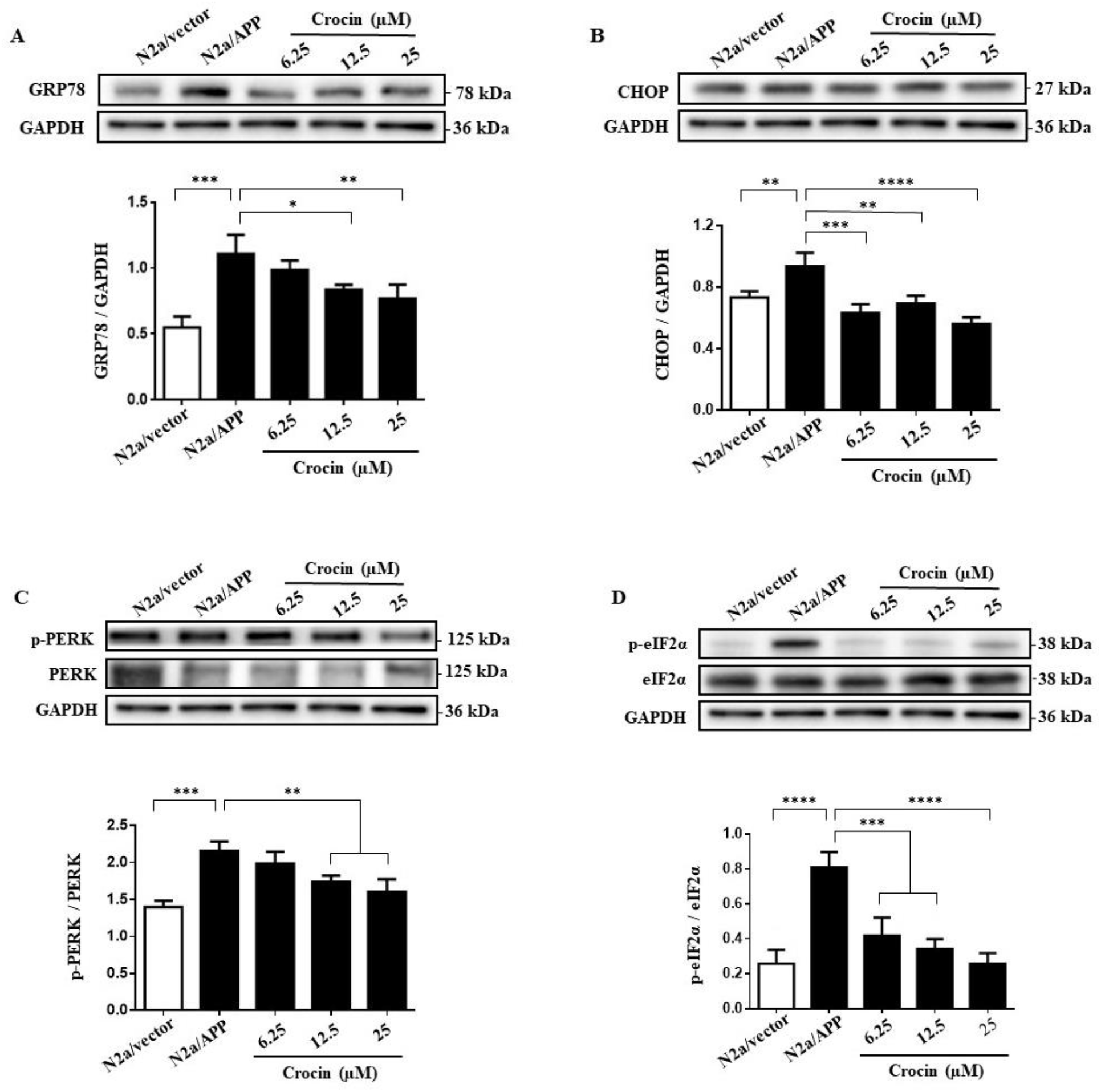
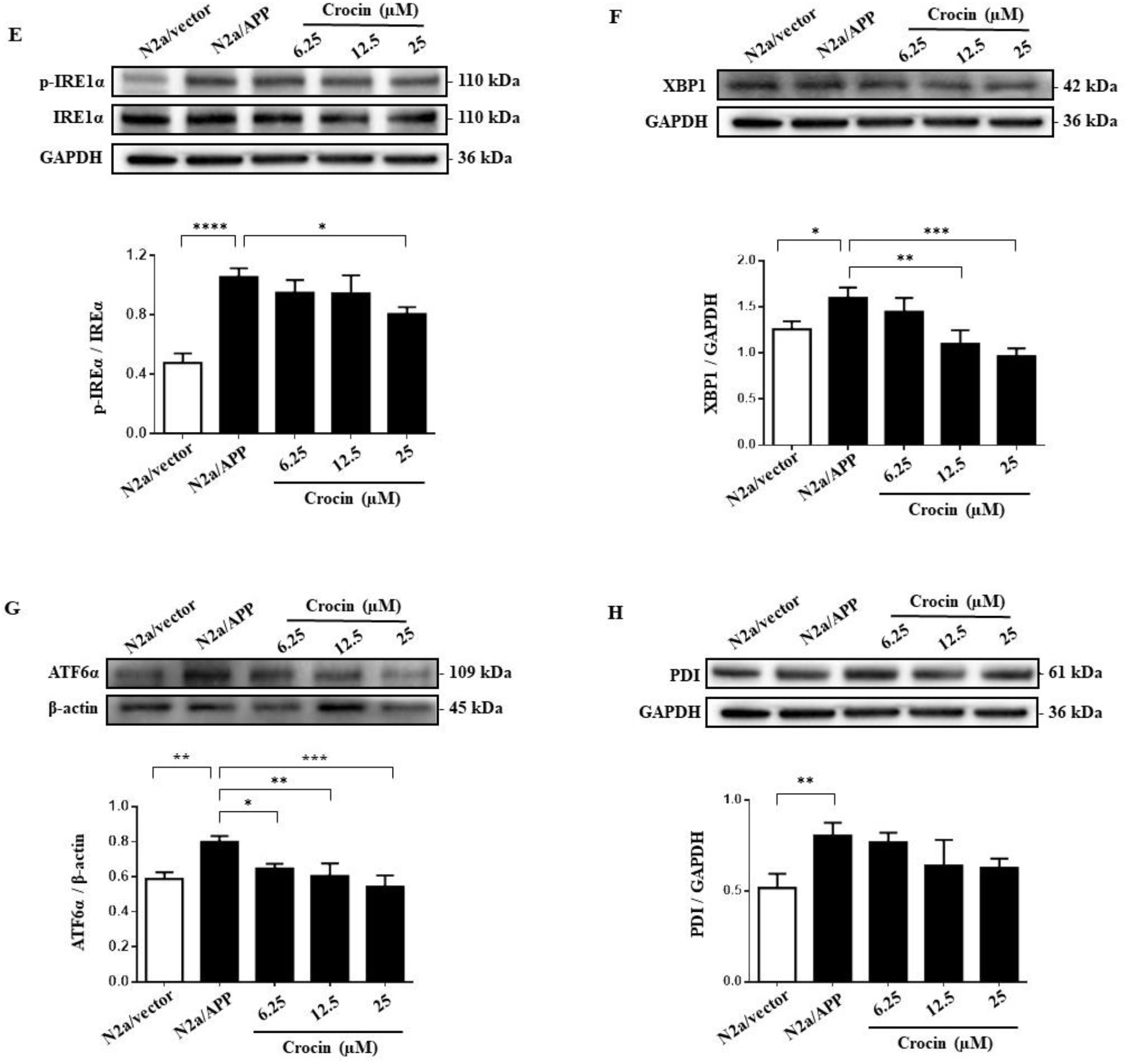
Crocin reduced ER stress of N2a/APP cells. (A-H) Proteins involved in the ER stress UPR pathways including GRP78, CHOP, p-PERK, p-eIF2α, p-IRE1, XBP1, ATF6α, and PDI in N2a/vector and N2a/APP cells were determined by Western blot analyses and the relative protein expression were shown in the bar charts, either GAPDH or β-actin was used as loading control. N = 3. \**p* < 0.05, \*\**p* < 0.01, \*\*\**p* < 0.001, \*\*\*\**p* < 0.0001 compared with N2a/APP cells treated with vehicle.

## Discussion

Recent studies have indicated that crocin could produce neuroprotective effects and may be beneficial for preventing cognitive dysfunction in AD. Crocin was effective at improving learning and memory in various mouse and rat models of AD, and reduced oxidative stress, Aβ deposition and cellular apoptosis in the brain [2-7]. Here, we demonstrated for the first time that crocin reduced Aβ generation through promoting α-cleavage of APP processing and inhibited ER stress by attenuating UPR signaling p-PERK/p-eIF2α and AT6α.

We showed that crocin significantly inhibited the protein expression of total APP in N2a/APP cells (Fig. 3B), and greatly promoted α-cleavage of APP processing to increase sAPPα generation (Fig. 3C) and reduce sAPPβ generation (Fig. 3D). On the other hand, we observed that crocin only modestly reduced the protein levels of BACE-1 and PS1 (a key subunit of the γ-secretase complex) (Fig. 3E-F). These results suggested that Aβ reduction by crocin was mainly associated with the non-amyloidogenic APP processing pathway. The relative proteolytic efficiency of the amyloidogenic and non-amyloidogenic APP processing is critical in Aβ peptide generation. Thus, our results showed that crocin was effective at blocking Aβ generation by decreasing APP expression and promoting α-cleavage of APP.

It is suggested that ER stress is associated with AD pathogenesis [14]. The ER is a primary intracellular site that misfolded proteins such as Aβ accumulate and initiate ER stress. It is reported that UPR signaling proteins including p-PERK, p-eIF2α and p-IRE1α are abnormally activated in the hippocampal neurons of AD patients [15, 16]. Increased levels of ER stress markers such as GRP78 (also known as BiP) and XBP1 were also detected in the brain cortex of 3xTg-AD mice [17]. Therefore, we investigated the effect of crocin on ER stress. We showed that ER stress and UPR proteins including GRP78, CHOP, p-PERK, p-eIF2α, p-IRE1α, XBP1, ATF6α and PDI were all significantly elevated in N2a/APP cells compared to N2a/vector. Our data showed crocin effectively reduced the levels of GRP78 and CHOP (Fig. 4A-B). The heat shock protein 70 (Hsp70) molecular chaperone GRP78 is an abundant ER-resident protein essential for many cellular processes. A major function of GRP78 in ER stress is to serve as a master regulator and regulate the activation of UPR signaling. When UPR activation is sustained, it may initiate apoptotic cell death via the up-regulation of the C/EBP homologous protein (CHOP). Thus, our data suggested crocin was effective at reducing ER stress at GRP78 level.

We further investigated the effects of crocin on the three key branches of UPR signaling pathways. Our data showed crocin was effective at inhibiting p-PERK/p-eIF2α, but inhibited p-IREα modestly. We also found that crocin significantly inhibited XBP1 and ATF6α. Under prolonged ER stress, the three branches of UPR pathways are abnormally activated. The autophosphorylation of PERK kinase activates downstream mediator p-eIF2α leading to inhibition of general protein translation and upregulation of ATF4 that induces the expressions of genes related to apoptosis [18, 19]. With the branch of IRE1α, IRE1α activation enables the splicing of XBP1 to XBP1s which upregulates the transcription of genes involved in ERAD and chaperones (e.g. GRP78, PDI) [19]. With the branch of AT6α, ER stress induces translocation of AT6α from ER to the Golgi, where it is cleaved and transferred into the nucleus to induce the expression of several UPR target genes (e.g. GRP78, CHOP, XBP1) [19]. Our data suggested crocin might be potential for preventing inhibition of protein synthesis and cellular apoptosis via attenuating p-PERK/p-eIF2α (Fig. 4C-D). The results also suggested crocin did not alter the upregulation of ERAD and ER chaperones transcription, which might be beneficial for Aβ degradation, as implicated by its minimal effects on p-IRE1α (Fig. 4E) and PDI expression (Fig. 4H). Interestingly, the level of XBP1 was significantly reduced by crocin (Fig. 4F), which might be explained by the effect of crocin on inhibiting AT6α (Fig. 4G).

In conclusion, the present study showed that crocin is able to promote α-cleavage of APP processing and reduce Aβ production. Crocin is also effective at inhibiting abnormal UPR activation induced by sustained ER stress induced observed in N2a/APP cells.

## Acknowledgements

Credit author statement: Crystal Cuijun Lin performed experiments and data analysis and drafted the manuscript. Erin Qian Yue contributed to experiment design and writing the manuscript. Zirong Liang contributed to writing the manuscript. Simon Ming Yuen Lee and Zaijun Zhang contributed to experimental design. Maggie Pui Man Hoi supervised and revised the manuscript. This work was funded by The Science and Technology Development Fund, Macau SAR (File no. 127/2014/A3, 0015/2019/ASC, 0023/2020/AFJ, 0035/2020/AGJ) and The University of Macau Research Grant (Project no. MYRG2015-0061-ICMS-QRCM, MYRG2017-00150-ICMS), and National Natural Science Foundation of China (NSFC) (Project No. 81403139).

